# Within host selection for faster replicating bacterial symbionts

**DOI:** 10.1101/222240

**Authors:** Ewa Chrostek, Luis Teixeira

## Abstract

*Wolbachia* is a widespread, intracellular symbiont of arthropods, able to induce reproductive distortions and antiviral protection in insects. *Wolbachia* can also be pathogenic, as is the case with *w*MelPop, a virulent variant of the endosymbiont of *Drosophila melanogaster*. An extensive genomic amplification of the 20kb region encompassing eight *Wolbachia* genes, called Octomom, is responsible for *w*MelPop virulence. The Octomom copy number in *w*MelPop can be highly variable between individual *D. melanogaster* flies, even when comparing siblings arising from a single female. Moreover, Octomom copy number can change rapidly between generations. These data suggest an intra-host variability in Octomom copy number between *Wolbachia* cells. Since *w*MelPop *Wolbachia* with different Octomom copy numbers grow at different rates, we hypothesized that selection could act on this intra-host variability. Here we tested if total Octomom copy number changes during the lifespan of individual *Drosophila* hosts, revealing selection for different *Wolbachia* populations. We performed a time course analysis of Octomom amplification in flies whose mothers were controlled for Octomom copy number. We show that despite the Octomom copy number being relatively stable it increases slightly throughout *D. melanogaster* adult life. This indicates that there is selection acting on the intra-host variation in the Octomom copy number over the lifespan of individual hosts. This within host selection for faster replicating bacterial symbionts may be in conflict with between host selection against highly pathogenic *Wolbachia*.

## Introduction

Gene copy number variation is one of the mechanisms allowing rapid evolution across the tree of life [1–3]. In bacteria, growth inhibition by nutrient limitation or antibiotic presence may be overcome by increasing copy number of genes functionally related to these challenges [4]. Moreover, amplified genomic regions allow accumulation of mutations without the risk of loss of the original function. This can lead to the generation of more beneficial variants and subsequent loss of extra copies or repurposing of the new copies for a new function [4]. Thus, genomic amplifications generate extensive and reversible genetic variation, which can either increase the fitness of an individual directly or be a substrate on which adaptive evolution can act.

We have previously found that a genomic amplification affects the biology of the intracellular, maternally transmitted bacterium *Wolbachia* [5]. *Wolbachia* is a widespread endosymbiont of insects, causing an array of phenotypes, including reproductive manipulations [6] and antiviral protection [7,8]. Moreover, some *Wolbachia* strains can strongly reduce the host lifespan. This was first described for *w*MelPop, a laboratory *Wolbachia* variant, in *Drosophila melanogaster* [9]. The Octomom genomic region, which contains eight *Wolbachia* genes, is amplified in *w*MelPop, while it is present as a single copy in closely related non-pathogenic *Wolbachia* variants [10,11]. The number of copies of this region varies greatly between individual *w*MelPop-infected flies from the same population, ranging from two to ten copies [5]. We have previously established *D. melanogaster* lines carrying defined and different Octomom copy numbers and observed that the higher the Octomom copy number, the higher *Wolbachia* levels and the shorter the lifespan of its *D. melanogaster* host [5]. Moreover, a *w*MelPop that reverted to carrying only one Octomom copy proliferates at the same rate as the control *w*MelCS_b variant and is not pathogenic [5]. Thus, we identified Octomom copy number as a pathogenicity determinant of *w*MelPop [5].

The high variation in Octomom copy numbers between individual flies can also be observed in the progeny of single *w*MelPop-carrying females [5]. This variation between siblings could be explained by *Wolbachia* variation within a female and differential symbiont assortment to the progeny. The fact that the variability decreased under selection argues for initial high variation within single flies, which is pruned over a few generations of selection for either the highest or the lowest Octomom copy number. However, even under constant selection some variation was either maintained or continuously generated, since reversing the direction of the selection or relaxing it could rapidly change the Octomom copy numbers in these lines [5].

We hypothesized that variation in Octomom copy number between *Wolbachia* cells within an individual host could lead to a differential growth of these cells. If *Wolbachia* cells with higher Octomom copy number proliferate more, their frequency in the pool of *Wolbachia* within a host will increase over time, and the average Octomom copy number of the within-host population increases over the host lifespan. We tested this hypothesis through a time course analysis of Octomom copy number in individual *w*MelPop flies originating from mothers with controlled Octomom copy number.

## Results

We examined the stability of Octomom copy number over the adult life (from eclosion to the onset of high mortality) in single *w*MelPop-carrying *D. melanogaster* (Fig 1A). The flies were the offspring of parents carrying *w*MelPop with low Octomom copy number (median of 4.5 Octomom copies, range from 4 to 5.5) or high Octomom copy number (median of 9.5, range from 8.5 to 11.5). The low Octomom cohort had, on average, 5.45 (standard deviation, SD = 0.82) Octomom copies per genome and the high Octomom cohort had an average of 10.29 (SD = 1.90) Octomom copies per genome. The fitted generalized additive model (GAM) clearly shows an increase in Octomom copy number in the first few days (six to eight days), which subsequently levels off at the later timepoints (Fig 1A and B). The trend is highly significant (the smooth term for time, *p* < 0.001). This shows that Octomom copy number in the *w*MelPop population increases during most of the adult host lifespan.

**Figure 1.**
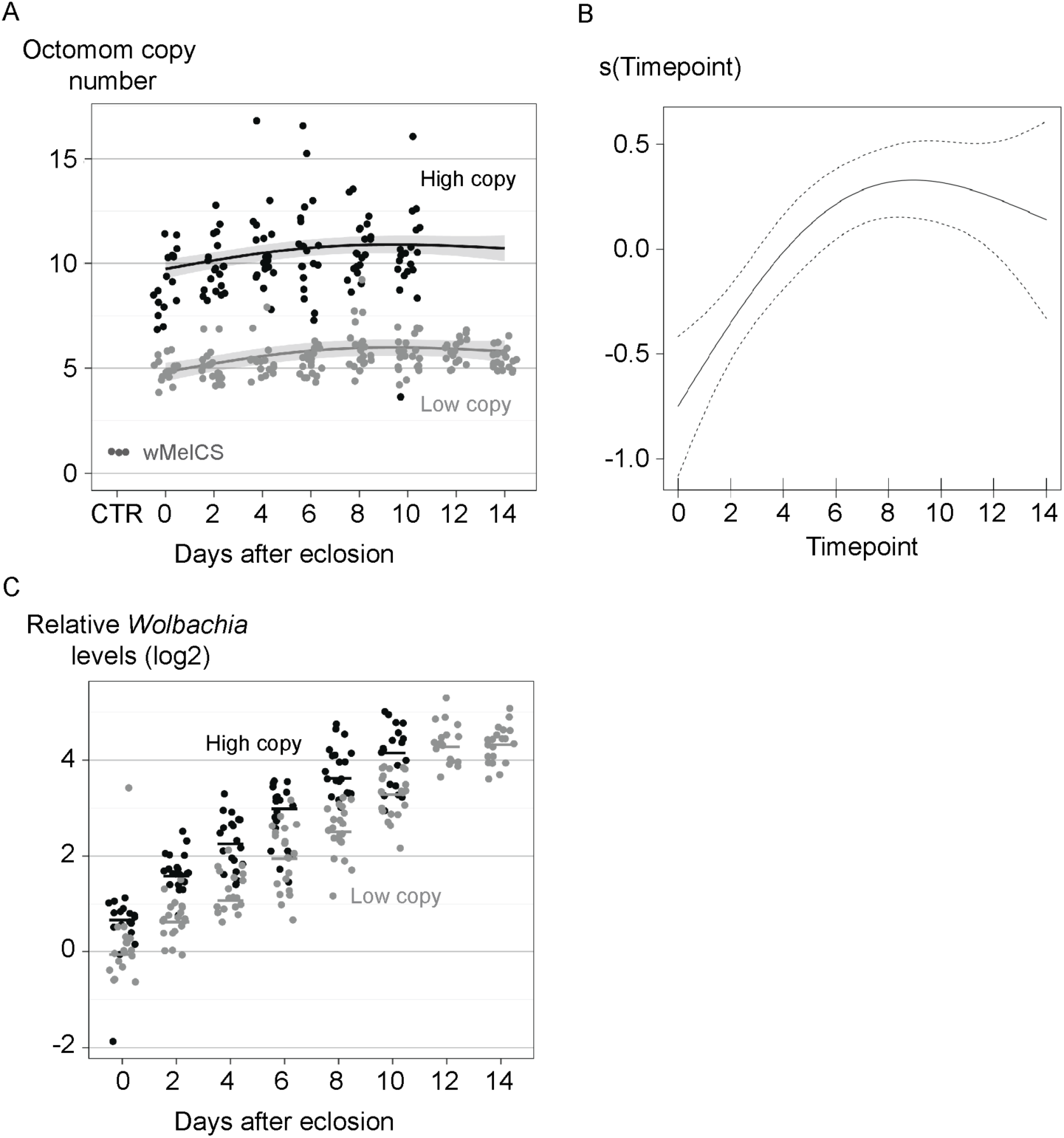
Octomom copy number and *Wolbachia* levels during adult *D. melanogaster* life. (A) Each dot represents *WD0513* genomic levels in a single fly, as we have previously shown that this gene can be used to estimate Octomom region copy number [5]. The values were obtained using the Pfaffl method with *wsp* as a reference gene and calibrated using the median of three samples of control *w*MelCS_b flies (CTR). The lines represent the fit of the generalized additive model (GAM) and shaded area - the 95% confidence interval. (B) Fitted GAM smooth of Octomom copy number in response to the age of adults. The 95% confidence interval is indicated by the dashed lines. Note the Y-axis is standardized so that average is zero. (C) Each dot represents *wsp* levels in a single fly. Lines are medians of the replicates. *wsp* levels were obtained using Pfaffl method with *Rpl32* as a reference gene and calibrated using the median of the low copy number samples at time zero. (A) and (C) represent data from single females derived from the 3^rd^ generation of selection for Octomom copy number in the DrosDel isogenic *w*^1118^ genetic background [5]. Females were raised and kept at 25°C (survival of their siblings is shown in Fig. S5A of [5]). Supporting data can be found in S1 Data.

In parallel, we tested *Wolbachia* titer in the same flies. *Wolbachia* levels increase with the age of flies (log-linear mixed-effect model (lme), *p* < 0.001) and flies carrying *w*MelPop with higher Octomom copy numbers have 1.85 times higher *Wolbachia* levels compared to flies carrying *w*MelPop with lower Octomom copy numbers (lme, *p* < 0.001), confirming previous results (Fig 1C). However, we do not see a significant interaction between cohort and growth rate (lme, *p* = 0.102). We also tested if Octomom copy number could be an explanatory variable for *Wolbachia* levels within each cohort. This variable was not significant (lme, *p* = 0.48).

## Discussion

By analyzing cohorts of individuals from mothers carrying *Wolbachia* with controlled Octomom copy number we observed that Octomom copy number is relatively stable over the life of the host. However, we detected a small, but clear and statistically significant, increase in Octomom copy number over time in the first six to eight days of the adult *D. melanogaster* life. This could be explained by selection acting on the heterogeneity of *w*MelPop copy number between *Wolbachia* cells within a single host. Since bacteria with higher copy numbers grow faster over time, they contribute more to the total pool of *w*MelPop in an older fly and therefore increase total Octomom copy number. In our dataset, the Octomom copy number stops increasing at the last days of the host life. This is surprising and could be explained by the initial heterogeneity in the flies being reduced in the course of selection. If *Wolbachia* cells with the maximal Octomom copy number within each fly reach a very high frequency or are fixed, there is no genetic variation for selection to act on, and the Octomom copy number does not continue to increase. Alternatively, the differential fitness between *Wolbachia* harboring different Octomom copy numbers may decrease with the age of the fly, weakening the strength of the selection and preventing continuous Octomom copy number growth.

Importantly, our data indicate that the selective pressures acting on *w*MelPop within and between hosts are different. Within a host, there may be a selection for *Wolbachia* that grow fast. On the other hand, competition between flies could select for *Wolbachia* that grow slower and have a lower cost for the host [5]. These opposing selective pressures may play a role in shaping the evolution of *Wolbachia* in natural populations.

Although the results presented here are compatible with our prediction of within host heterogeneity and selection, other forces may contribute to the changes in Octomom copy number during host lifespan. Amplification and deletion of Octomom copies may also occur throughout *w*MelPop host’s adult life. The rates of these two opposing mutations will determine an overall tendency of Octomom copy number to increase or decrease over time. Therefore, the dynamics of Octomom copy number changes may be the combination of selection and mutation. A mutation bias for deletion of Octomom copies could temper the effect of selection for higher Octomom copy numbers. On the other hand, a bias for amplification could explain the observed increase in Octomom copy number, even in the absence of selection. Nonetheless, this would also result in an increased proportion of more pathogenic *w*MelPop with host age. The influence of mutation and selection on Octomom copy number change during host lifespan may vary with intrinsic and extrinsic factors. For instance, mutation bias towards amplification or deletion could change with Octomom copy number itself. On the other hand, since temperature has a strong effect on *w*MelPop pathogenicity [9,12], it may also affect the differential growth of *Wolbachia* with different Octomom copy numbers and, therefore, within host selection on this trait. A mechanistic understanding of how Octomom copy number changes and influences *Wolbachia* phenotype and which factors modulate it will help disentangling the relative contribution of selection and mutation to Octomom copy number changes during *D. melanogaster* life.

*Wolbachia* with higher Octomom copy number proliferate more and are more pathogenic to their hosts [5]. The within host selection for bacteria that proliferate more and have a higher potential of being deleterious to the host may be a common phenomenon. For instance, *Staphylococcus aureus* variants that cause blood or deep tissue infection are, in the majority, the result of within-host selection from non-pathogenic nose colonizing variants [13]. The adaptations conferring high virulence identified in this study do not favor *S. aureus* dissemination and onward transmission (discussed in [13]). Thus, the selective pressure acting on the bacteria within a single host may be in conflict with the selective pressure acting on the entire bacterial population. This implies that although the more pathogenic bacterial variants arise throughout the life of the hosts, they are constantly purged from the overall bacterial population by selection either on the fitness of the host (vertically transmitted symbionts) or on the bacterial transmission capacity (horizontally transmitted symbionts).

A recent report suggested that *w*MelPop Octomom copy numbers change drastically during *D. melanogaster* lifespan, increasing more than two-fold in the first ten days of adult life and then decreasing more than four-fold over the next thirty days [14]. These results differ from the ones we present here and may be explained by the lack of experimental control for the Octomom copy number in these flies and determination of the Octomom copy numbers after the onset of mortality. Using data from Chrostek and Teixeira 2015 [5] we constructed a model showing that in a mixed population of flies the differential growth of *Wolbachia* with different Octomom copy numbers, combined with differential death of flies carrying *Wolbachia* with different Octomom copy numbers, leads to initial increase in Octomom copy number, followed by a decrease due to death of the flies carrying *Wolbachia* with higher Octomom copy number at the host population level [15].

Here we confirmed that *w*MelPop with higher Octomom copy number has higher *Wolbachia* titers, supporting our conclusion that this amplification controls *w*MelPop levels. However, the difference in growth rate between these lines was not statistically significant. This may be due to the high variability in the data and the differential growth between these lines being potentially small. We have previously shown different growth rates between *w*MelPop carrying one and two copies of Octomom and between these and *w*MelPop carrying 12 or 15 copies [5]. However, the growth rate of *w*MelPop carrying 12 and 15 copies was not significantly different [5]. This indicates that the relationship between growth rate and Octomom copy number is not linear and that differences in *w*MelPop carrying higher copy numbers may have a smaller impact on growth. Therefore, the difference in growth between *w*MelPop carrying five and ten copies may indeed be small. The difference in *Wolbachia* titers despite the lack of measurable difference in the growth rate might be the result of the cumulative effect of small growth rate differences throughout the fly development from egg to adult, the result of a differential growth rate at different development stages, or even the accumulation of small differences in growth for more than one generation.

The labile nature of the Octomom amplification and the resulting phenotypes of this amplification make *w*MelPop an interesting case study to understand genome dynamics and selective forces acting on endosymbionts. This system may be further used in the future to reveal general principles in host-bacteria symbiosis.

## Material and methods

### Fly strains

*D. melanogaster* DrosDel isogenic background (*iso*) flies with *w*MelCS_b and *w*MelPop were described before [5,10]. Selection on *D. melanogaster* lines carrying *w*MelPop with different Octomom copy number was described in [5].

### Experimental setup for time-course analysis of *WD0513* and *Wolbachia* levels

Female progeny of females from the 3^rd^ generation of selection for Octomom copy number in the *D. melanogaster* DrosDel isogenic background (*iso*) [5] was collected at eclosion (ten females per tube), allowed to mate with brothers for 24 h (five males per tube), separated from males, and 20 females were sacrificed every second day for *WD0513* and *Wolbachia* density quantification. Females were maintained at 25°C on a standard cornmeal diet without live yeast and were passed to fresh vials every 3 days. We sampled only until the onset of high mortality in the different lines in order to avoid sampling bias for surviving, low Octomom copy number bearing flies.

### DNA extractions

DNA was extracted from single flies (*w*MelPop) or pools of ten flies (*w*MelCS_b controls). Each fly or pool of flies was squashed in 250 μl of 0.1 M Tris HCl, 0.1 M EDTA, and 1% SDS (pH 9.0) and incubated 30 min at 70°C. Next, 35 μl of 8 M CH_3_CO_2_K was added, and samples were mixed by shaking and incubated on ice for 30 min. Subsequently, samples were centrifuged for 15 min at 13,000 rpm at 4°C, and the supernatant was diluted 100× for qPCR.

### Real-time quantitative PCR

The real-time qPCR reactions were performed using CFX384 Real-Time PCR Detection System (Bio-Rad) as described before [5,10]. Each reaction contained 6 μl of iQ SYBR Green Supermix (Bio-Rad), 0.5 μl of each primer (3.6 mM), and 5 μl of diluted DNA. We performed two technical replicates for each sample for each set of primers. Primer sequences were described before [5]. For all three genes assayed: *Wolbachia WD0513* and *wsp*, and *Drosophila Rpl32* the following thermal cycling protocol was applied: 2 min at 50°C, 10 min at 95°C, and 40 cycles of 30 s at 95°C, 1 min at 59°C, and 30 s at 72°C. Melting curves were examined to confirm the specificity of amplified products. Ct values were obtained using Bio-Rad CFX Manager software with default threshold settings. Ct values were subjected to a quality check - samples with standard deviation between technical replicates exceeding 0.5 for one of the genes were discarded. The experiment spanned six qPCR plates and three samples of ten wMelCS_b flies (extracted and aliquoted beforehand and assayed on every qPCR plate) were used to normalize between plates. Relative amounts of genes were calculated by the Pfaffl method [16]. To apply the method, the efficiency of each primer set was predetermined in a separate experiment. For relative Octomom copy number quantification, *WD0513* was the target gene and the single-copy *wsp* gene was used as a reference. The medians of three samples of pools of ten wMelCS_b flies were used as control values for the Pfaffl method. *w*MelCS_b has one copy of the Octomom region in the genome, determined by the coverage analysis of sequencing data [10]. This sample, with known Octomom copy number, is required to estimate Octomom copy number of the remaining samples [17]. For *Wolbachia* quantification, *wsp* was the target gene and *Drosophila Rpl32* gene was used as a reference. The levels of *wsp* are relative to the median of the samples of the low Octomom cohort at time zero.

### Statistical analysis

The statistical analysis was performed in R [18]. The script of the analysis is provided in S1 Text. Graphs were generated using the package ggplot2 [19].

Since the temporal trend over time of the number of Octomom copies was not linear we analyse it by fitting a Generalized Additive Model (GAM, package mgcv in R [20]). We included time and line as independent variables and PCR plate as a random effect. The smooth terms for the interaction between time and lines were nonsignificant (*p* > 0.114) and were removed form the final model.

Analysis of *wsp* levels over time was performed with log-linear mixed-effect model fits (package lme4 in R [21]). The effect of interaction between factors was determined by an ANOVA comparing models fit to the data with and without the interaction.

## Financial Disclosure

LT lab is funded by the Fundação para a Ciência e Tecnologia (www.fct.pt) grant PTDC/BEX-GMG/3128/2014. EC is supported by EMBO Long Term Fellowship EMBO ALTF 1497-2015 (http://www.embo.org), co-funded by Marie Curie Actions by the European Commission (LTFCOFUND2013, GA-2013-609409) (ec.europa.eu/research/mariecurieactions). The funders had no role in study design, data collection and analysis, decision to publish, or preparation of the manuscript.

## Acknowledgments

We thank Tiago Marques for advice on the statistical analysis.

## Supporting information

S1 Data – Relative levels of *WD0513* and *wsp* in single females carrying wMelPop.

S1 Text – R script with the statistical analysis of the data.

